# Laccase-mediated catalyzed fluorescent reporter deposition for live cell imaging

**DOI:** 10.1101/731539

**Authors:** Brandon Cisneros, Neal K. Devaraj

## Abstract

Catalyzed reporter deposition (CARD) is a widely established method for labeling biological samples analyzed using microscopy. Horseradish peroxidase, commonly used in CARD to amplify reporter signals, requires the addition of hydrogen peroxide which may perturb samples used in live-cell microscopy. Herein we describe an alternative method of performing CARD using a laccase enzyme, which does not require exogenous hydrogen peroxide. Laccase is an oxidative enzyme which can carry out single-electron oxidations of phenols and related compounds by reducing molecular oxygen. We demonstrate proof-of-concept for this technique through the non-targeted covalent labeling of bovine serum albumin using a fluorescently-labeled ferulic acid derivative as the laccase reporter substrate. We further demonstrate the viability of this approach by performing live-cell CARD with an antibody-conjugated laccase against a surface bound target. CARD using laccase produces an amplified fluorescence signal by labeling cells without the need for exogenous hydrogen peroxide.

---

Laccases (EC 1.10.3.2, benzenediol:oxygen oxidoreductase) are a family of enzymes which couple the reduction of molecular oxygen to the oxidation of a substrate, typically a phenol or aromatic amine. Laccases carry out four one-electron oxidations of its substrate and use these electrons to reduce oxygen to water. This process is mediated by a cluster of four copper atoms in its active site. Laccases are found in many organisms, notably fungi[1] and plants, but also in insects[2] and bacteria,[3] and they show variable substrate specificity across species.

Owing to the versatile nature of the chemical transformation they carry out, laccases have been adopted or proposed for use in many industrial processes.[4] Much of laccase’s utility in industry has focused on decolorizing dye[5] or treating wood pulp used to produce paper.[6] Moreover, laccase has been suggested as a useful agent for treating contaminated water,[7] and removing phenol constituents in various food or beverages[8]. In academic research, laccase has been used for the chemical transformations of small molecules,[9] production of polymers,[10] as an enzyme in ELISA,[11–13] and in biosensing applications.[14] This research has led to the development of modified laccases with a variety of properties, including stability in organic solvents[15] and improved reactivity in biological media.[16]

Catalyzed reporter deposition (CARD) is a technique that is frequently used in the preparation of tissues or cells for microscopy.[17] In this technique, an analyte-dependent reporter enzyme is used to deposit an easily-detectable reporter molecule near the desired target. This is often achieved by modifying the enzyme with an affinity tag, typically by linking it to an antibody, and using it to chemically activate a biotin-labeled or fluorophore-labeled substrate.[18,19] The activated substrate can then covalently bind to the targeted cell (Figure 1A).[17] Enzyme turnover allows for many more fluorescent labels to be deposited per antibody delivered compared to an antibody that has been directly labeled with a fluorescent tag. This approach has been widely used with horseradish peroxidase (HRP) but has also been used with other enzymes such as alkaline phosphatase.[19]

**Figure 1.**
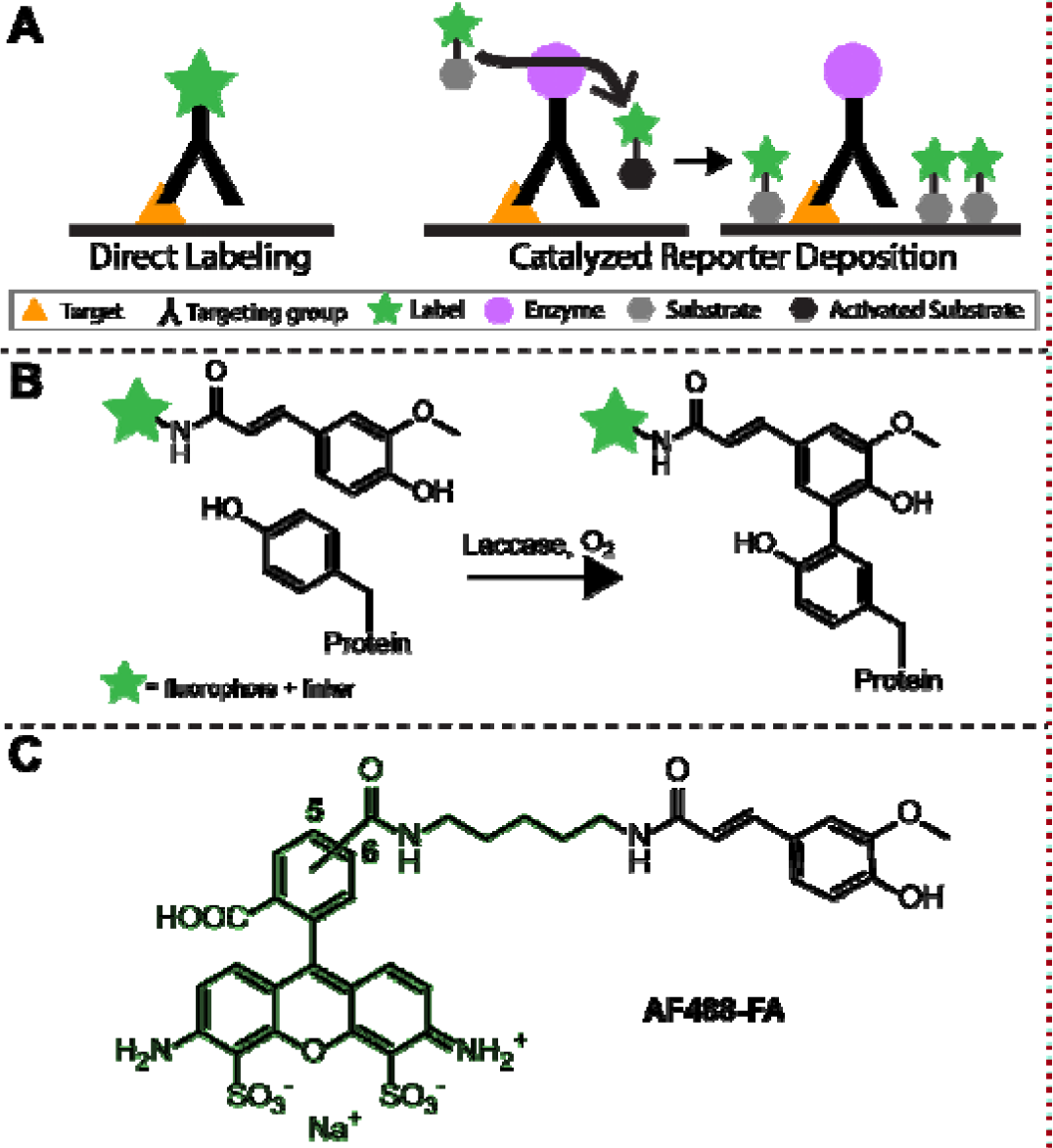
A) Graphical depiction comparing direct labeling to CARD. The fluorescent label is covalently attached to the targeting group in direct labeling, whereas the fluorescent label is covalently deposited in the area near the target in CARD. B) Schematic representation of the way by which AF488-FA could covalently bind to tyrosine residues in a protein. C) Structure of AF488-FA which consists of ferulic acid bound to Alexa Fluor 488 via a cadaverine linker.

More recently, genetically encoded engineered ascorbate peroxidase (APEX) has been used for amplified imaging in cells for fluorescent microscopy,[20] electron microscopy,[21] and proteomic labeling.[22] Unlike typical CARD methodologies, APEX uses a genetically encoded reporter enzyme as a label. The addition of heme during cell culture allows for reconstitution of enzyme activity that can subsequently be used to deposit a probe (e.g. biotin-phenol). Using reporter enzymes to provide an amplified signal for imaging and labeling has proven to be a useful tool in studying biological systems in many different contexts.[18,22–26]

One caveat of using a peroxidase such as HRP or APEX is that these techniques use exogenous hydrogen peroxide as the oxidizing agent. Hydrogen peroxide has been shown to have a number of effects on cells at various concentrations, including inducing differentiation, proliferation,[27] DNA lesions,[28] and apoptosis.[29] Even at lower concentrations, hydrogen peroxide is recognized to alter cell signaling,[30] particularly in sensitive cells like neurons.[31] These effects are highly dependent on the specific conditions and identity of the cells in question.[32] The diversity of effects over a range of concentrations makes it difficult to predict what might happen upon addition of hydrogen peroxide to any given cell line. As a result, CARD with peroxidases can have limitations when used for live cell imaging.

We hypothesized that laccase could be used to carry out CARD for live cell fluorescence microscopy imaging. This seemed plausible because laccase and HRP can both oxidize various phenolic substrates and carry out similar chemical transformations of their substrates albeit via distinct mechanisms.[33–35] Since laccase uses oxygen as the oxidizing agent and produces water, cells would not be exposed to unnecessarily levels of hydrogen peroxide required by HRP. Because laccases oxidize their substrates to products that are identical to those produced by peroxidases, these products are expected to react with tyrosine residues to form adducts (Figure 1B).[36]

To test whether laccase could serve as a suitable enzyme for carrying out CARD, we used two commercially available laccases derived from different organisms: Chinese lacquer tree (Toxicodendron vernicifluum formerly Rhus vernicifera) and turkey tail mushroom (Trametes versicolor). Initial attempts at verifying previously reported enzyme activities and pH optima using syringaldazine[37–39] were difficult to reproduce, possibly due to the heterogenous quality of the enzyme preparations. Reproducibility was considerably improved by triturating the enzyme powder with a minimum quantity of buffer in a hand homogenizer before adding the remaining quantity of buffer. This suspension was then syringe filtered to remove large particles and concentrated via spin filtration to produce a transparent, stable solution of laccase.

To assess the viability of using laccase to perform CARD, we selected ferulic acid as a suitable substrate due to its high rate of reaction with our chosen laccases.[39] We designed a probe (Figure 1C) consisting of Alexa Fluor 488 conjugated to ferulic acid via a cadaverine linker. The probe was synthesized by reaction of succinimidyl ferulate with Alexa Fluor 488 cadaverine and subsequently purified by semi-preparative HPLC. The resulting probe (AF488-FA) was dissolved in dimethyl sulfoxide and was used in all subsequent CARD tests.

We tested the ability for laccase to perform CARD first in an untargeted fashion (Figure 2A). By co-incubating laccase with the probe and a soluble protein target, bovine serum albumin (BSA), we expected that some of the probe should be oxidized to the ferulic acid semiquinone. The oxidized probe could then react with tyrosine residues on the protein target, leading to conjugation. After performing such an incubation, the solutions were subjected to repeated washing and spin dialysis. Washes included both denaturing and ethanolic conditions to ensure that non-covalently bound AF488-FA was removed. Both laccases were found to be able to modify BSA in solution but T. versicolor laccase provided better conjugation yields at pH 7.0 150 mM phosphate buffer, 19.5 μM AF488-FA, and 0.5% (w/v) BSA (Figure 2B). Variable concentrations of BSA showed that increasing concentrations of BSA do not noticeably increase non-specific binding (Figure 2C).

**Figure 2.**
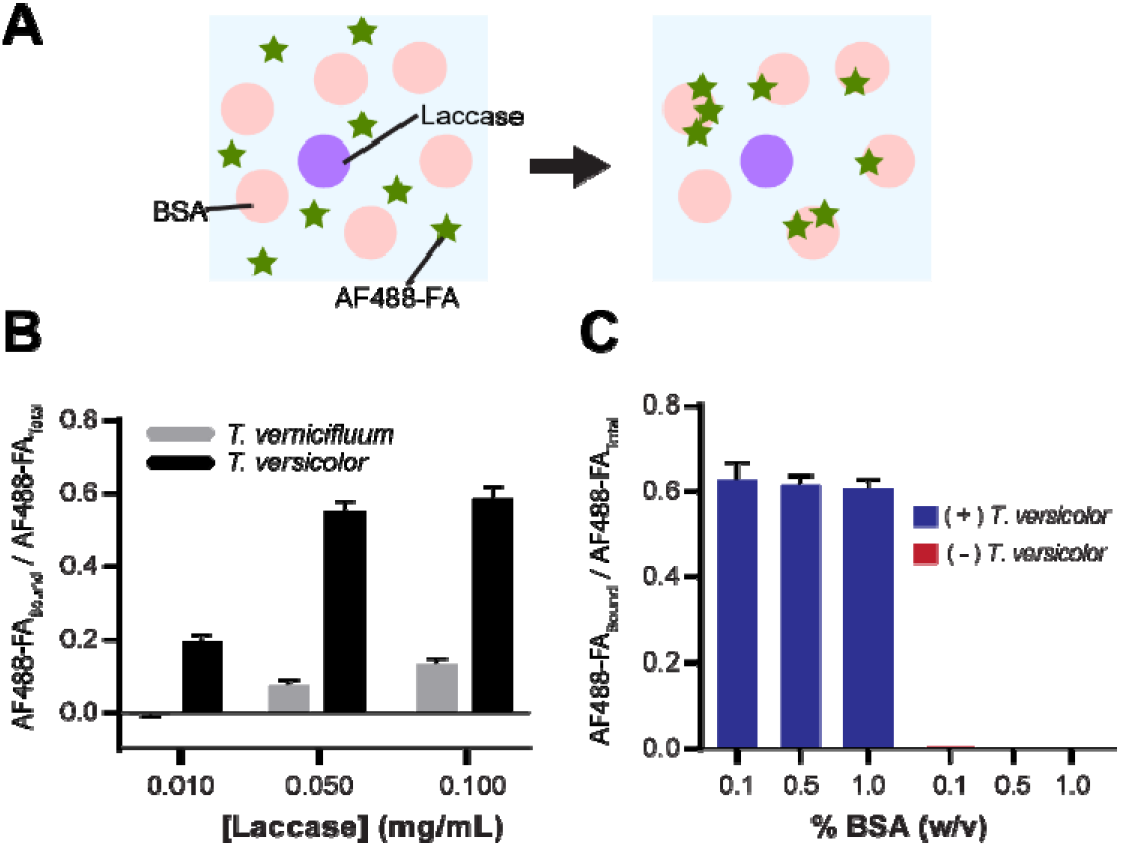
A) Representation of non-targeted labeling of BSA with AF488-FA using laccase. BSA in the general proximity of a laccase can be covalently labeled when activated AF488-FA diffuses close to it and reacts with amino acid residues (e.g. tyrosine) in its structure. B) Fraction of AF488-FA bound to BSA after washing with various concentrations of laccase showing superior performance of T. versicolor at all enzyme concentrations. Refer to Supplementary Information for detailed conditions. C) Dependence of BSA modification on BSA concentration.

We set out to determine if laccase could be used to fluorescently label the surface of living cells with AF488-FA via CARD. We labeled cells with a biotinylated primary antibody followed by a laccase-streptavidin conjugate (laccase-SA) (Figure 3A). Our approach mimicked the use of a primary antibody followed by an enzyme-conjugated secondary antibody for fluorescent labeling of fixed cells as in CARD. The modular nature of this approach could allow the laccase-SA to be used for other biotinylated antibodies or targets. In addition, an antibody that has laccase attached to it could be easily prepared by pre-incubating the laccase-SA with a biotinylated antibody.

**Figure 3.**
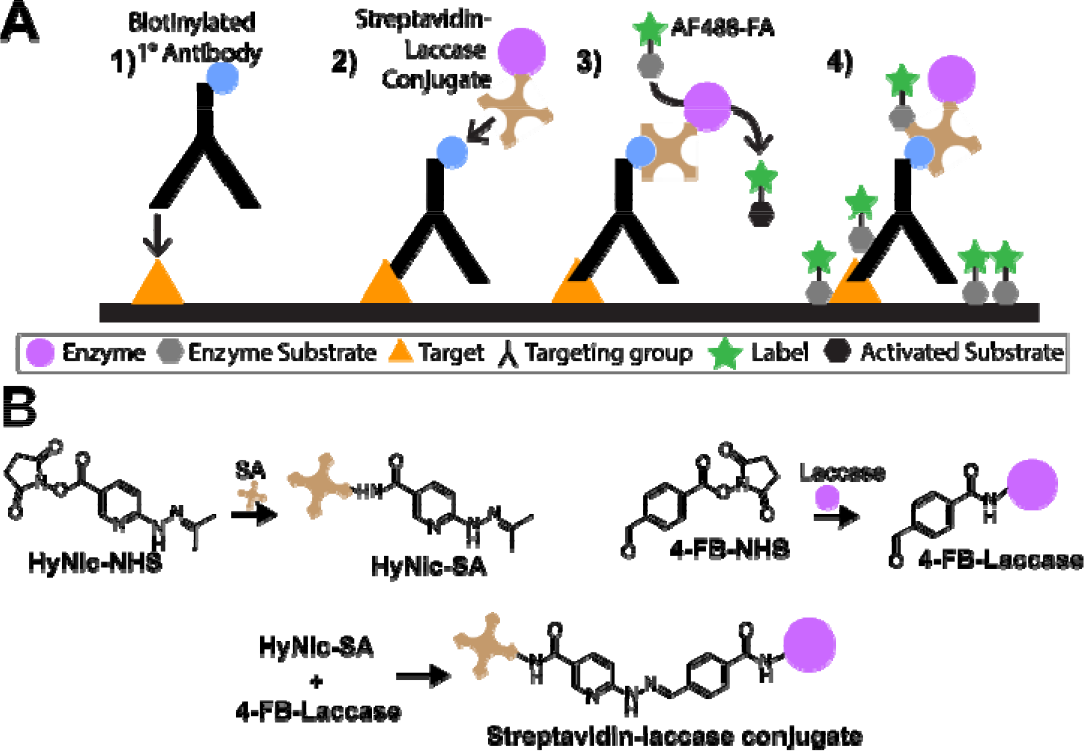
A) A biotinylated primary antibody can interact with a target on the surface of a cell (1). The streptavidin moiety of a laccase-SA can bind to biotin on the primary antibody (2). Exposure to the AF488-FA probe will cause the probe to become activated by the laccase (3). The activated probe can react with amino acids on the cell surface proteins nearby (4). B) Synthesis of HyNic-modified streptavidin, 4-FB-modified laccase, and streptavidin-laccase conjugate.

To create the laccase-SA, streptavidin and laccase were modified with amine-reactive succinimidyl 6-hydrazinonicotinate acetone hydrazone (HyNic-NHS) and succinimidyl 4-formylbenzoate (4-FB-NHS), respectively, using a Solulink protein-modification kit. The derivatized proteins were subsequently reacted with one another to yield the streptavidin-laccase conjugate. The resulting product was found to have 1.1 laccases per streptavidin molecule based on the optical detection of the hydrazone formed between HyNic and 4-FB. Conjugation was further validated by measuring laccase enzymatic activity using syringaldazine[40] as a colorimetric substrate.

To test laccase CARD on live cells, we chose to target epidermal growth factor receptor (EGFR) on the surface of A431 cells. A431 is a human squamous cell carcinoma cell line that has high levels of surface-expressed EGFR.[41] EGFR has been intensively studied as a therapeutic and imaging target, and has been used as a model system for surface receptor internalization and recycling.[42] For our imaging experiments, we grew A431 cells in 8-well chamber slides. Prior to imaging, the cells were sequentially incubated with the biotinylated anti-EGFR primary antibody at 4 °C for 30 min, laccase-SA for 15 min at room temperature, and the AF488-FA in phosphate buffer at 37 °C for 30 min. The low temperatures were used to minimize receptor internalization.[43] As a comparison, cells were also treated with commercial Alexa Fluor 488-labeled streptavidin (SA-488) (4 fluorophores/protein) using the same duration, temperature, and concentration of streptavidin as was used for the streptavidin-laccase conjugate. The confocal fluorescence microscopy images of these cells indicate substantial signal amplification of the laccase-SA and AF488-FA treated cells compared to the Alexa Fluor 488-streptavidin conjugate (Figure 4A and Figure S3).

**Figure 4.**
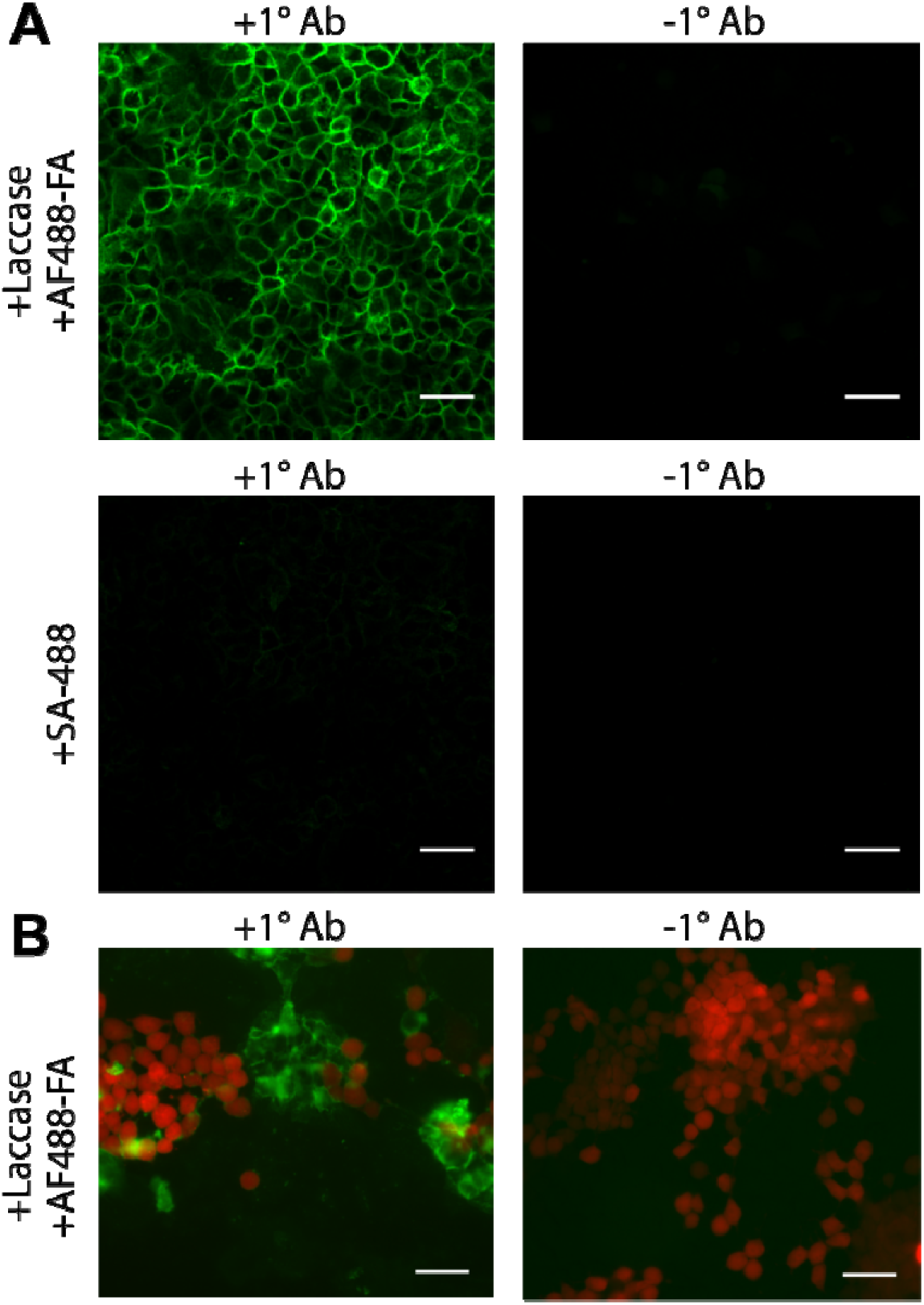
A) Confocal microscopy images of A431 cells treated with biotinylated anti-EGFR primary antibody followed by either laccase-SA/AF488-FA or SA-488. Control cells (“−1° Ab”) were subjected to the same conditions but without the primary antibody. Scale bar = 50 μm. B) Epifluorescence microscopy images of merged AF488 and mCherry channels of co-culture of A431 cells and mCherry-expressing HEK293 cells. Scale bar = 50 μm All microscopy images here are window-leveled identically.

We set out to quantify the amplified fluorescence signal produced by this method for a population of cells. Cells were grown in a black-bottomed 96-well culture plate and were fluorescently labeled with laccase/AF488-FA or SA-488. The cells were fluorescently labeled using the same methods as were used for microscopy. The total fluorescence for each well was measured on a plate reader (Figure S4). These data also indicate that the total fluorescence was greater for the laccase-SA treated cells than for the SA-488 treated cells. This indicates that the quantity of fluorophores deposited per cell and thus the total fluorescence was amplified relative to the stoichiometric labeling method. These results could be readily improved with further research and optimization of conditions such as concentration of AF488-FA, duration of incubation, or identity of the fluorescent substrate.

To qualitatively assess the specificity of this approach in a mixed cell population, we labeled and imaged co-cultured cells. A431 cells (+EGFR) and mCherry-expressing HEK293 cells (−EGFR) were grown together in chamber slides and were prepared in the same manner as other live-cell imaging experiments. The epifluorescence microscopy images obtained (Figure 4B and Figure S5) show that the EGFR-expressing A431 cells (−mCherry) show high levels of probe deposition by AF488-FA whereas the EGFR-deficient HEK293 cells (+mCherry) do not. This demonstrates that CARD with laccase can selectively label live cells expressing a target of interest in mixed cell populations.

We then tested whether CARD with laccase changes the uptake of epidermal growth factor (Figure S6). A431 cells were labeled with laccase, as before, but were subsequently treated with 20 ng/μL Alexa Fluor 555 EGF Complex (EGF-AF555) for 1h at 4 °C to allow EGF-AF555 to bind. Cells were then incubated at 37 °C for 30 minutes to elicit endocytosis.[44] Confocal microscopy of these cells shows that the overall signal is diminished in cells treated with the anti-EGFR antibody compared to those without. Puncta are observed in both cases, but this suggests that the primary anti-EGFR antibody used for CARD with laccase reduces the overall intensity of EGF-AF555 signal observed. As a result, CARD with laccase could plausibly inhibit EGF signaling to some degree. This observation is likely caused by a combination of two factors: direct antibody interference with the EGF binding site on EGFR and receptor endocytosis during laccase labeling.[45] Optimization of the amount of primary antibody needed could potentially improve EGF receptor/ligand binding.

We have demonstrated that laccase can be used to fluorescently label proteins via CARD. Antibody-targeted laccases enable biomarker dependent amplification of imaging signals. We selectively labeled live cells with AF488-FA and imaged them without fixation, even in the presence of other cell types. This technique allows for signal amplification in a manner akin to APEX or CARD with horseradish peroxidase, but it does not require exogenous hydrogen peroxide. With further evaluation of other laccases and laccase substrates, this technique could be modified to accept different reporter substrates and have improved reactivity, optimal pH, ion sensitivity, etc. We envision that laccase-based CARD techniques could be potentially useful for many labeling applications and could serve as a basis for an improved method of performing amplified reporter deposition of live cells in biomarker fluorescence imaging, electron microscopy, and proteomics.

## Acknowledgements

This work was funded by the NIH under grant number DP2DK111801. Author B.T.C. acknowledges support from the National Science Foundation Graduate Research Fellowship under Grant No (DGE-1650112)

