## Supplemental Information for "Laccase-mediated catalyzed fluorescent reporter deposition for live cell imaging"

B. T. Cisneros, and N. K. Devaraj

Abstract: Catalyzed reporter deposition (CARD) is a widely established method for labeling biological samples analyzed using microscopy. Horseradish peroxidase, commonly used in CARD to amplify reporter signals, requires the addition of hydrogen peroxide which may perturb samples used in live-cell microscopy. Herein we describe an alternative method of performing CARD using a laccase enzyme, which does not require exogenous hydrogen peroxide. Laccase is an oxidative enzyme which can carry out single-electron oxidations of phenols and related compounds by reducing molecular oxygen. We demonstrate proof-of-concept for this technique through the non-targeted covalent labeling of bovine serum albumin using a fluorescently-labeled ferulic acid derivative as the laccase reporter substrate. We further demonstrate the viability of this approach by performing live-cell CARD with an antibody-conjugated laccase against a surface bound target. CARD using laccase produces an amplified fluorescence signal by labeling cells without the need for exogenous hydrogen peroxide.

**Experimental Procedures**

**1. Materials and Instrument Details**

Laccases (Trametes versicolor and Toxicodendron vernicifluum), syringaldazine, dicylcohexylcarbodiimide solution, and THF purchased from Sigma Aldrich (St. Louis, MO). trans-Ferulic acid was purchased from Tokyo Chemical Inc. (Tokyo, Japan). All other reagents and solvents were obtained from Fisher Scientific. Solulink Protein-Protein Conjugation kit (S-9010) was purchased from TriLink Biotechnologies (San Diego, CA). Biotinylated anti-EGFR antibody clone 528 (#MA5-12872), AlexaFluor 488-cadaverine, streptavidin-AlexaFluor 488, and AlexaFluor 555 EGF Complex (EGF-AF555) were purchased Thermo Fisher (Waltham, MA). Fetal bovine serum (FBS) was purchased from Corning (Corning, NY). All unspecified products (e.g. buffer components, etc.) were purchased from Fisher Scientific (Waltham, MA). pHR_Gal4UAS_tBFP_PGK_mCherry[1] was a gift from Wendell Lim (Addgene plasmid # 79130). pLP1 (gal-pol) and pLP2 (rev) were obtained from Invitrogen (Waltham, MA). pCMV-G was obtained from the Friedmann lab.[2]

UV-Vis spectroscopy for BCA assay, fluorophore concentration determination, and kinetics assays were performed on a Nandrop 2000 (Thermo Fisher, Waltham, MA).

Reverse-phase HPLC purification was performed using an Agilent 1100 series Infinity HPLC (Santa Clara, CA). Agilent Zorbax SB-C18 semi-prep column (ID 9.4 x 250 mm, 5 µm, 80 Å) using a water/acetonitrile gradient containing 0.1% trifluoroacetic acid using a variable wavelength detector wavelength of 495 nm. High resolution mass spectroscopy was collected on an Agilent Infinity 1260 LC and tandem Agilent 6230 high resolution time of flight (TOF) mass spectrometer managed by the UCSD Department of Chemistry and Biochemistry Molecular Mass Spectroscopy Facility.

Confocal fluorescence microscopy imaging was performed on an Axio Observer Z1 motorized inverted microscope (Carl Zeiss Microscopy GmbH, Germany) with Yokogawa CSU-X1 spinning disk confocal unit to and an Evolve 512x512 EMCCD camera (Photometrics, Canada) using ZEN imaging software (Carl Zeiss Microscopy GmbH, Germany). Fluorophores were excited with laser diodes (488 nm; 30 mW). Epifluorescence microscopy imaging for coculture experiments was performed on an Axio Observer D1 inverted microscope (Carl Zeiss Microscopy, GmbH, Germany) with an Axiocam MRm camera (Carl Zeiss, Microscopy, GmbH, Germany) and an HBO-100 mercury arc lamp epifluorescence source (Carl Zeiss Microscopy, GmbH, Germany). A 49008 ET mCH/TR filter cube (Chroma Technology Group, Bellows Falls, VT) and 38 HE filter cube (Carl Zeiss Microscopy, GmbH, Germany) were used for imaging of fluorophores. Images were processed using Image J[3] with Fiji.[4] Microscopy images shown in a given figure are window-leveled in the same manner as one another.

**2. Synthesis of N-succinimidyl ferulate (NHS-FA)**

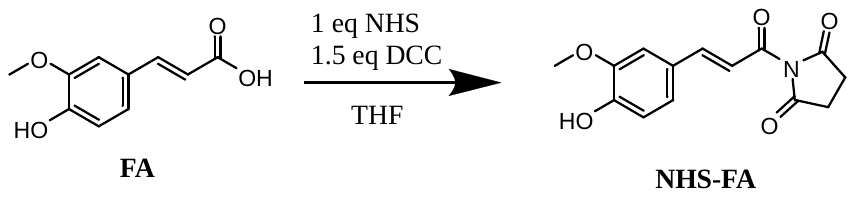

Scheme S1: Synthesis of N-succinimidyl ferulate

250 mg of ferulic acid (1.27 mmol, 1 eq.) and 146 mg (1.26 mmol, 1 eq.) of NHS were dissolved in 5 mL tetrahydrofuran. This solution was chilled over ice and 1.91 mL (1.91 mmol, 1.5 eq.) of 1 M dicyclohexyl carbodiimide (DCC) was added to the reaction. The reaction was monitored with TLC (1:1 hexane:ethyl acetate). After 24 hours, the reaction filtered over a medium frit and rotary evaporated to dryness. The residue was suspended in methylene chloride causing formation of a white precipitate. The solution was filtered through a short plug of silica and purified by column chromatography (1:1 hexane:ethyl acetate).

**3. Synthesis of AlexaFluor 488-cadaverine-ferulate (AF488-FA)**

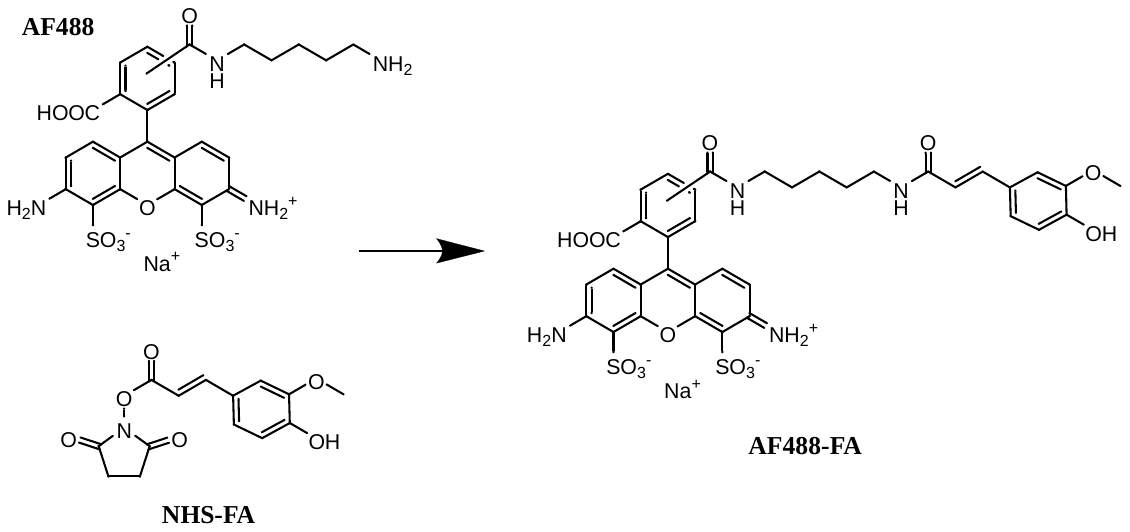

Scheme S2: Synthesis of AlexaFluor 488-cadaverine-ferulate (AF488-FA)

Solutions of AlexaFluor 488-cadaverine and NHS-FA in DMF were prepared at 2.0 mg/mL and 1.0 mg/mL, respectively. 100 μL (0.31 μmol) of the AlexaFluor 488 cadaverine solution, 180 μL (0.65 μmol, ~2 eq.) of the NHS-FA solution, and 8 μL (0.046 μmol, 0.15 eq.) of diisopropylethylamine were combined and incubated at 45 °C for 8 hours while shaking. The solution was concentrated in vacuo, dissolved in acetonitrile, and purified by HPLC. Recovered yield: 0.13 mg (53%) as determined by UV-Vis spectroscopy. HRMS (ESI-TOF) calculated for [C36 H33 N4 O13 S2]- ([M ]- ) 793.1491, found 793.1484 (Supplemental Figure S1)

Table S1. HPLC gradient conditions for purification of A488-FA

| Time[a] | Water + 0.1% TFA | Acetonitrile + 0.1% TFA |
| --- | --- | --- |
| 0 min | 95% | 5% |
| 5 min | 80% | 20% |
| 12 min | 70% | 30% |
| 15 min | 70% | 30% |

[a] The product eluted at approximately 12-12.4 min.

**4. High-Resolution Time-of-Flight Mass Spectrometry of AF488-FA**

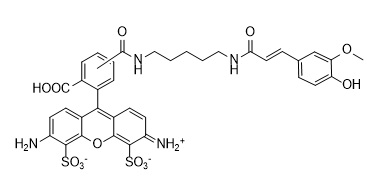

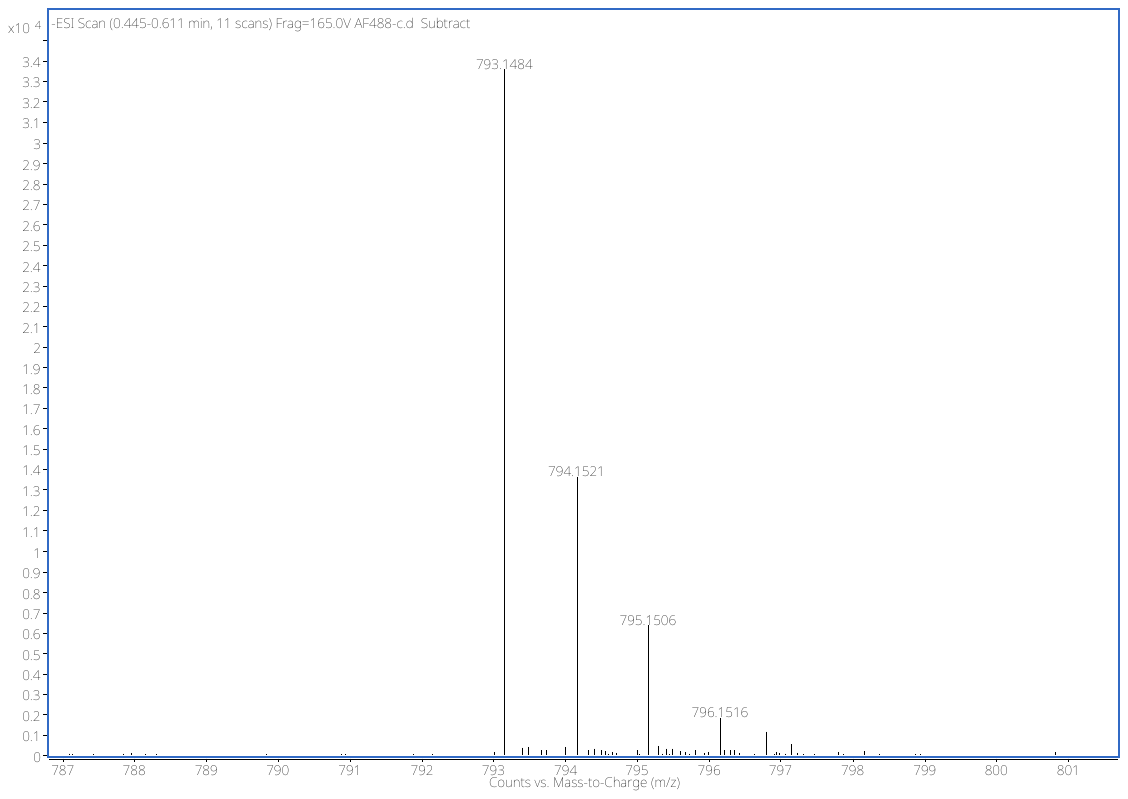

[M]^-^

Figure S1: HRMS spectrum of AF488-FA

Table S2. HRMS results data

| Mass Measured | Theoretical Mass | Delta (ppm) | Composition |
| --- | --- | --- | --- |
| 793.1484 | 793.1491 | -0.9 | [C36H33N4O13S2]- |

**5. Laccase Resuspension**

Laccase was brought into solution prior to modification and conjugation. To suspend, laccase was triturated with warm (~37 °C) buffer (typically pH 7.0 150 mM sodium phosphate buffer) in a Dounce homogenizer. Buffer was added in small portions and removed, being careful to not pipet any enzyme powder that remains undissolved. This was continued until all buffer had been thoroughly triturated with the enzyme. The resulting suspensions were dark and turbid with some visible sediment. These were incubated at 37 °C for 1 hour with gentle shaking. After incubating, the samples were centrifuged at 5,000 rcf for 5 minutes. The supernatant was syringe filtered with a 0.22 micron filter to remove large particles. The resulting transparent, light brown/yellow solution was concentrated with a centrifuge filter (e.g. Amicon Ultra 10 kDa or similar) to produce a dark yellow-brown solution that lacked turbidity. This solution was stable for 1 month at 4 °C without significant loss of activity.

Protein concentration of these solutions were determined with BCA assay and enzymatic activity was determined with a syringaldazine assay.

**6. Syringaldazine Laccase Activity Assay**

A 0.216 mM solution of syringaldazine was prepared in methanol. Laccase was suspended in an appropriate buffer at the desired pH. 2.2 mL of buffer and 0.5 mL of laccase solution were added to a UV-Vis cuvette with a stir bar and the solution was allowed to reach 37 °C in a UV-Vis spectrometer. Upon equilibration, 0.3 mL of the 0.216 mM syringaldazine solution (pre-warmed to 37 °C) was added and A530 was measured every 5 seconds for 10 minutes. From the linear portion of the resulting graph, units of laccase activity was calculated as follows:

$$\frac{Units}{mL} enzyme=\frac{\Delta A_{530}}{0.001 \times0.5 mL}$$

This same protocol can be adapted to a microscale format where measurements of 2 microliter samples are taken at longer time intervals on the pedestal of the Nandrop instead of in a cuvette. The above equation can be used slight alteration. Care should be taken to use 1 cm absorbance measurements instead of 1 mm absorbance measurements in this case.

**7. Kinetic data of T. versicolor and T. vernicifluum**

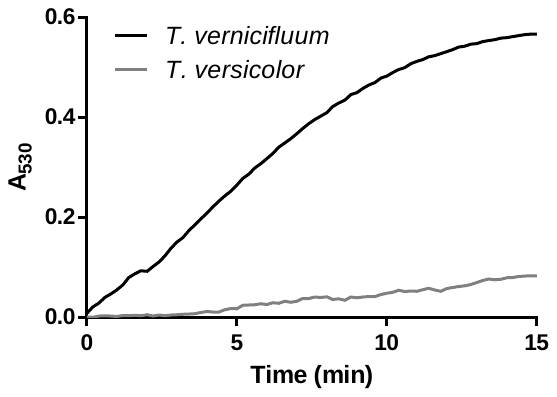

Figure S2: Representative kinetic curves of T. vernicifluum and T. versicolor at final concentration of 0.0167 mg/mL in 3 mL pH 7.0 150 mM sodium phosphate buffer with 10% MeOH, and 0.0216 mM syringaldazine at 37 °C.

If too many units of laccase are added to this assay, the absorbance from the colored product will begin to disappear over the observation scale of a kinetics experiment as the product is likely further oxidized. In this case, the amount of enzyme added to the assay and it should be decreased and the assay repeated.

**8. Non-targeted BSA Labeling Assay**

Variable enzyme concentration experimental conditions:

Incubation conditions: pH 7.0 phosphate buffer, 37°C, 90 min, 19.5 µM AF488-FA, 0.5% (w/v) BSA. The sample was then concentrated and subsequently washed in a 30k MWCO spin filter with: 1 × 8M urea, 1 × 50%(v/v) EtOH in pH 7.0 phosphate buffer, 5 × pH 7.0 phosphate buffer). Concentration of bound and unbound AF488-FA was determined spectrophotometrically using UV-Vis.

Variable BSA experimental conditions:

Same as above but a fixed concentration of enzyme at 0.1 mg/mL final concentration.

**9. Protein-Protein Conjugation**

A slightly modified version of the SoluLink protein conjugation protocol was followed. In brief:

The proteins to be modified were suspended/buffer exchanged into 250 mM sodium phosphate pH 8.0 and protein concentration was determined with BCA assay. The protein was concentrated to a >2 mg/mL for optimal reaction. The manufacturer-provided protein-conjugation excel spreadsheet was used to determine the amount of the S-4FB and S-HyNic reagents to add using a 10x excess of the succinimidyl ester. S-HyNic was reacted with laccase and S-4FB was reacted with streptavidin. They were incubated for 4 hours.

After reaction, the proteins were desalted using a buffer exchange column into pH 6 150 mM sodium phosphate. Protein concentration was determined using BCA assay and degree of modification was assessed by UV-Vis (as per manufacturer instructions). Modified laccase and streptavidin were combined at a 1:2 ratio and incubated at room temperature overnight. It is critical to not add the TurboLink catalyst. This catalyst is aniline based and will act as a substrate to laccase under the reaction conditions causing formation of a dark polymeric precipitate.

Resulting product was subjected to spin filtration with a 100kD MWCO to eliminate unconjugated laccase. UV-Vis was used to estimate degree of labeling of the proteins. Protein concentration was determined with BCA assay. Activity per mg of protein was determined with the syringaldazine laccase assay to further validate the degree of labeling determined by UV-Vis.

**10. Cell Culture**

A431, HEK293, and HEK293T cells (American Type Culture Collection), were cultured in DMEM with glutamine and pyruvate (GIBCO Thermo Fisher, Waltham, MA) 10% FBS (Corning, Corning, NY). Cells were detached for passaging using TrypLE Express (Thermo Fisher, Waltham, MA). LabTekII 8-well chamber slides were purchased from Nunc (Thermo Fisher, Waltham, MA).

**11. Lentiviral Production and Transduction**

HIV1-based lentivirus vectors were produced by transient co-transfection of HEK293T cells. HEK293T cells in ten 150 mm dishes were co-transfected by polyethyleneimine (PEI) method with pHR_Gal4UAS_tBFP_PGK_mCherry, pLP1 (gal-pol), pLP2 (rev), and pCMV-G1. 12 ug of pHR_Gal4UAS-tBFP_PGK_mCherry, 12 μg pLP1, 10 μg pLP2, 6 μg pCMV-G, and 90 μg PEI were used per 150 mm plate. Conditioned media at day 1, 2, and 3 post-transfection were collected, filtered through a 0.45 micron filter, and concentrated by centrifugation at 7000 rpm for 16 hours at 4 °C with a Sorvall GS-3 rotor. The resulting pellets were resuspended with buffer containing 10 mM Tris HCl, pH 7.8, 1 mM MgCl2, and 3% (w/v) sucrose.

HEK293 cells were transduced with the resulting lentivirus by adding 30 microliters of concentrated virus to the HEK293 cells that were grown in a single well of a 24-well plate with 1 mL of DMEM with 10% FBS. The medium was changed after 2 days. Fluorescence revealed mCherry expression was sufficient for co-culture experiments.

**12. Live-Cell Microscopy Protocol**

Cells were grown in 8 well chamber slides (Nunc LabTek II) to a desired confluency (~70-90%). Culture medium was removed carefully by gently aspirating it from the corner of a slide with a pipette while tilting the slide. Being gentle is important as to not disrupt the monolayer of cells. Cells were incubated with biotinylated anti-EGFR antibody at a concentration of 0.002 mg/mL in whole media (including FBS) at 4 °C for 30 min. Longer incubation times can be used to improve labeling, but the kinetics of EGFR receptor internalization do not allow for particularly long labeling times (e.g. overnight at 37 °C). The cells are then washed three times by gentling rocking them in room temperature Ca- and Mg-free Hank’s Balanced Salt Solution. The supernatant is aspirated and replaced with either streptavidin-AlexaFluor 488 (SA-488) or the laccase-streptavidin conjugate.

The laccase-streptavidin conjugate was added to each well to a final concentration of 0.002 mg/mL. SA-488 was added to a final concentration of 0.00084 mg/mL. This concentration corresponds to an equivalent concentration of streptavidin as the laccase-streptavidin. These were incubated at room temperature for 15 min and then the cells were washed as before. For SA-488-treated cells, the media was replaced with pH 7 phosphate buffer. For laccase-streptavidin treated cells, the media was replaced with AF488-FA at 19.5 μM in pH 7 phosphate buffer. The samples were then incubated for 30 min at 37 °C. They were then washed as above and pH 7.4 phosphate buffered saline was added. The cells were then promptly imaged.

For fluorescent labeling experiments with EGF-AF555, after labeling cells with laccase, cells are washed with ice cold PBS and 20 ng/uL EGF-AF555 in cold 0.3% (w/v) BSA in DMEM and cells were incubated for 1hr at 4 °C. Cells were then incubated at 37 °C for 30 minutes to promote receptor endocytosis.

**Results and Discussion**

**13. Microscopy of Laccase-Streptavidin with AF488-FA and Streptavidin-Alexa Fluor 488**

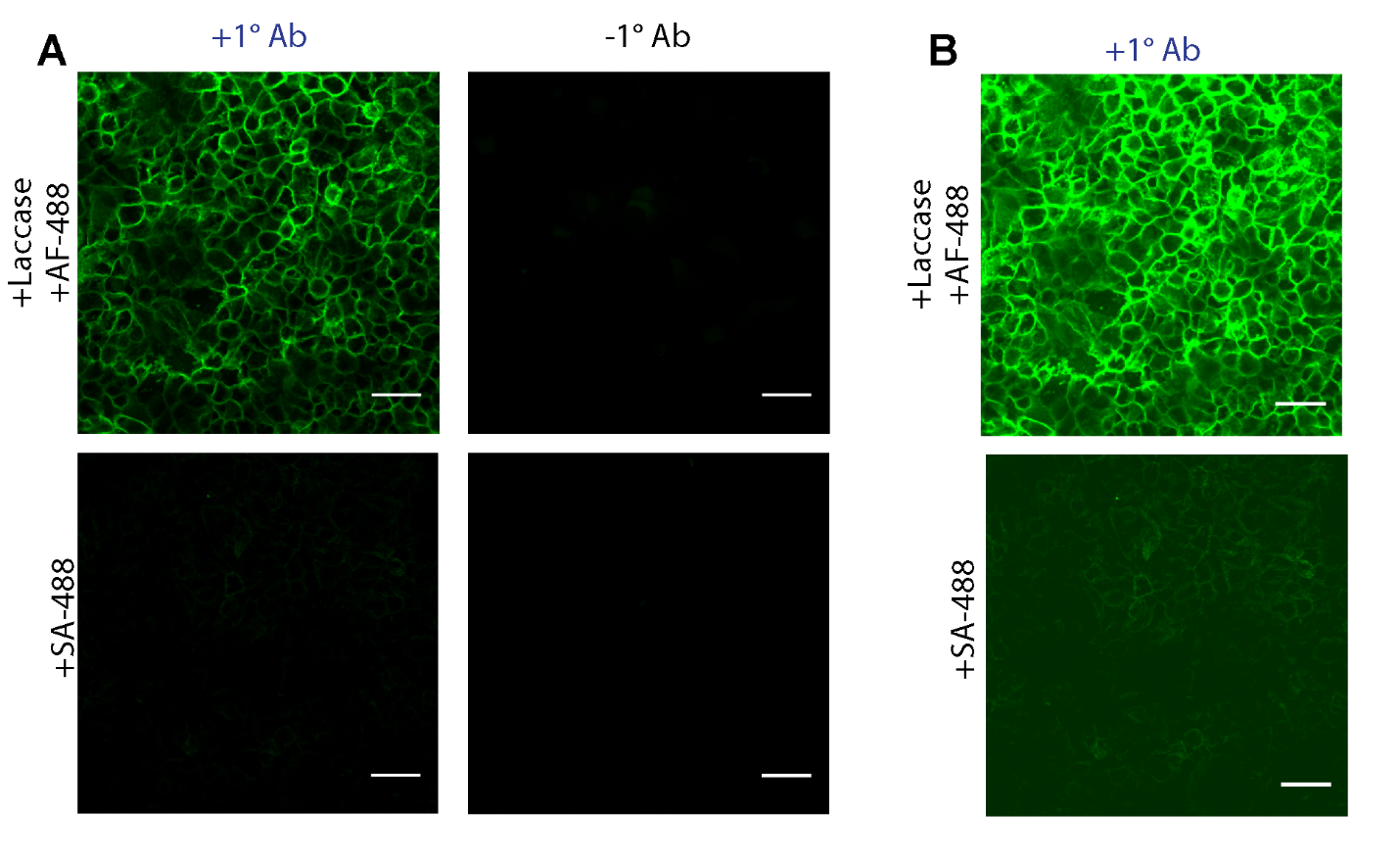

Figure S3: A) Fluorescence confocal microscopy images of signal amplification from probe deposition by laccase-streptavidin as compared to streptavidin-AlexaFluor 488. Images are taken under the same conditions and window-leveled identically. B) Images brightened to show similar pattern of fluorescence labeling for both SA-488-labeled and AF488-FA. Scale bar = 50 μm.

**14. Plate reader assay of Laccase/AF-488 and SA-488 cells**

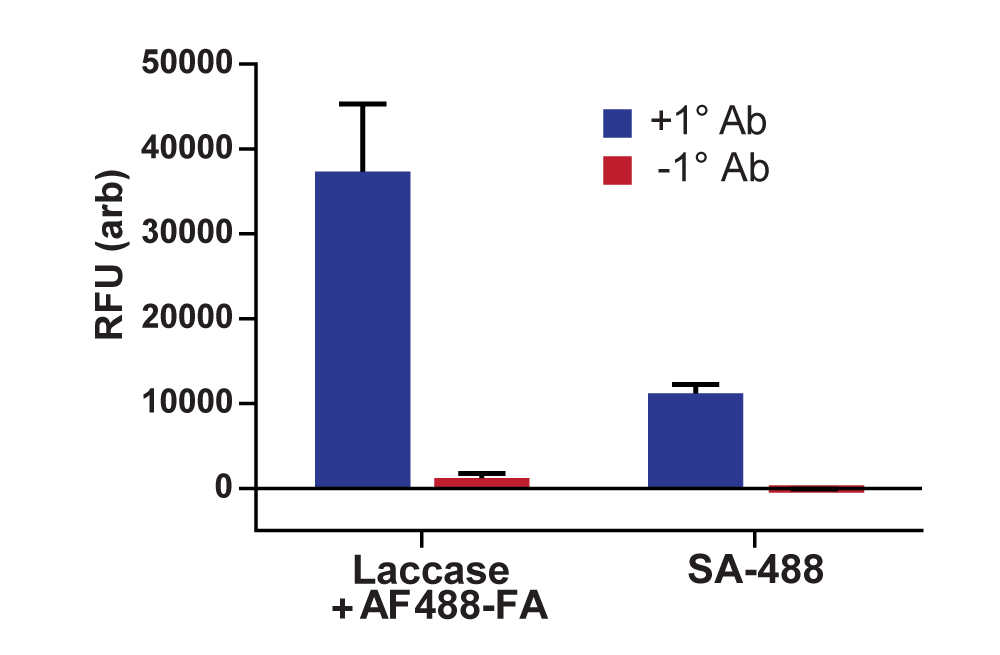

Figure S4: Bulk fluorescence measurements of cells obtained with a fluorescence plate reader, after being treated by the same method used for microscopy.

**15. Representative Co-Culture Microscopy Images**

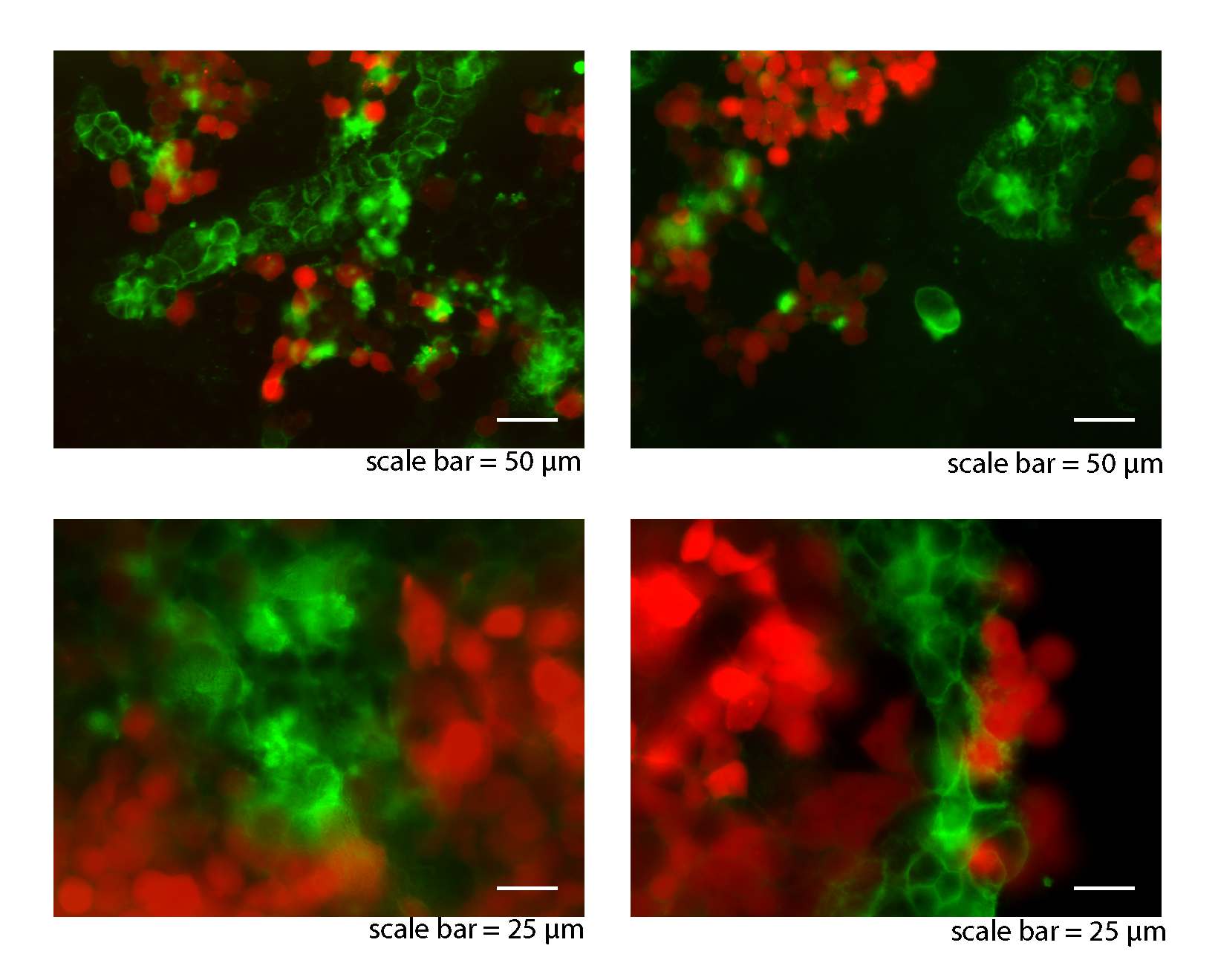

Figure S5: More example representative images of mCherry-expressing HEK293 (red) and AF488-FA labelled A431 cells (green). The top row of images are at 20x magnification and the lower row are at 40x magnification. All fluorescent images were acquired with epifluorescence microscopy.

**15. Fluorescent Epidermal Growth Factor Internalization Microscopy Images**

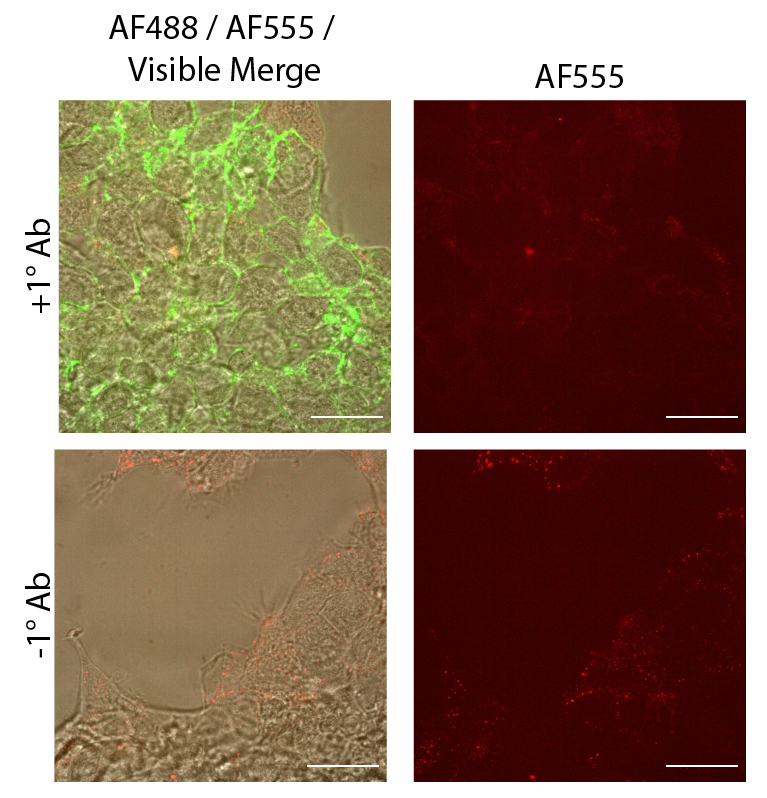

Figure S6: Confocal microscopy images of A431 cells treated with biotinylated anti-EGFR primary antibody followed sequentially by laccase-streptavidin, AF488-FA, and EGF-Alexa Fluor 555 conjugate. Cells were incubated for 30 minutes at 37 °C to elicit endocytosis. Control cells (“-1° Ab”) were subjected to the same conditions but without the primary antibody. Scale bar = 25 µm. All microscopy images here are window-leveled identically.

**Author Contributions**

Author B. T. Cisneros carried out the experimental design, data collection, analysis, and wrote the manuscript.

Author N. K. Deveraj was responsible for funding acquisition, assisted in experimental design, and assisted in editing the draft.
